# Efficient Generative Modelling of Protein Structure Fragments using a Deep Markov Model

**DOI:** 10.1101/2021.06.22.449406

**Authors:** Christian B. Thygesen, Ahmad Salim Al-Sibahi, Christian S. Steenmanns, Lys S. Moreta, Anders B. Sørensen, Thomas Hamelryck

**Author notes:** Correspondence to: Christian B. Thygesen < >, Thomas Hamelryck < >. Equal contribution (Anders B. Sørensen, Thomas Hamelryck).

## Abstract

Fragment libraries are often used in protein structure prediction, simulation and design as a means to significantly reduce the vast conformational search space. Current state-of-the-art methods for fragment library generation do not properly account for aleatory and epistemic uncertainty, respectively due to the dynamic nature of proteins and experimental errors in protein structures. Additionally, they typically rely on information that is not generally or readily available, such as homologous sequences, related protein structures and other complementary information. To address these issues, we developed BIFROST, a novel take on the fragment library problem based on a Deep Markov Model architecture combined with directional statistics for angular degrees of freedom, implemented in the deep probabilistic programming language Pyro. BIFROST is a probabilistic, generative model of the protein backbone dihedral angles conditioned solely on the amino acid sequence. BIFROST generates fragment libraries with a quality on par with current state-of-the-art methods at a fraction of the run-time, while requiring considerably less information and allowing efficient evaluation of probabilities.

## 1. Introduction

Fragment libraries (Jones & Thirup, 1986) find wide application in protein structure prediction, simulation, design and experimental determination (Trevizani et al., 2017; Chikenji et al., 2006; Boomsma et al., 2012). Predicting the fold of a protein requires evaluating a conformational space that is too vast for brute-force sampling to be feasible (Levinthal, 1969). Fragment libraries are used in a divide-and-conquer approach, whereby a full length protein is divided into a manageable sub-set of shorter stretches of amino acids for which backbone conformations are sampled. Typically, sampling is done using a finite set of fragments derived from experimentally determined protein structures. Fragment libraries are used in state-of-the-art protein structure prediction frameworks such as Rosetta (Rohl et al., 2004), I-TASSER (Roy et al., 2010), and AlphaFold (Senior et al., 2019).

Generally, knowledge-based methods for protein structure prediction follow two main strategies: homology (or template-based) modelling (Eswar et al., 2006; Šali & Blundell, 1993; Song et al., 2013) and *de novo* modelling (Rohl et al., 2004). Both approaches assume that the native fold of a protein corresponds to the minimum of a physical energy function and make use of statistics derived from a database of known proteins structures (Alford et al., 2017; Leaver-Fay et al., 2013). Whereas homology modelling relies on the availability of similar structures to limit the search space, knowledge-based *de novo* protocols require extensive sampling of the conformational space of backbone angles (figure 1).

**Figure 1.**
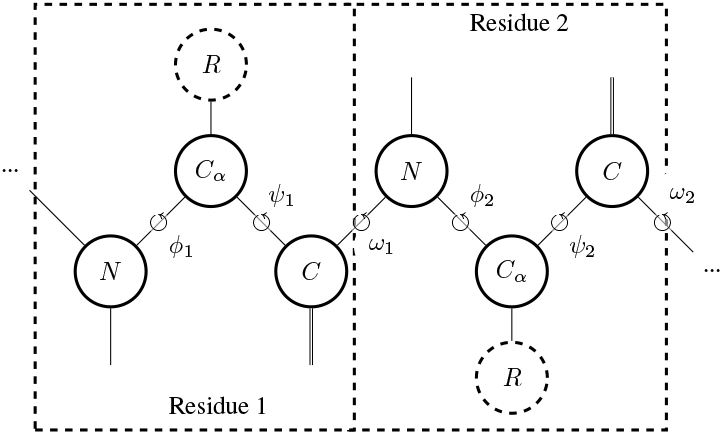
Schematic of the three dihedral angles (*ϕ, ψ*, and *ω*) that parameterise the protein backbone. *R* represents the side chain.

To overcome the shortcomings of either strategy, modelling tools like Rosetta (Rohl et al., 2004) use a combined approach of extensive sampling and prior information. Rosetta employs simulated annealing of backbone conformations according to an energy function (Alford et al., 2017), while reducing the conformational space by sampling fragments of typically 3 or 9 amino acids at a time (Simons et al., 1997).

Fragments are typically extracted from experimentally determined protein structures in the Protein Data Bank (Berman et al., 2000) and used in prediction based on similarities in sequence and sequence-derived features (Gront et al., 2011; Kalev & Habeck, 2011; Santos et al., 2015; De Oliveira et al., 2015; Trevizani et al., 2017; Wang et al., 2019). Generative probabilistic models of protein backbone angles (Hamelryck et al., 2006; Boomsma et al., 2008; Bhattacharya et al., 2016; Edgoose et al., 1998; Li et al., 2008; Lennox et al., 2010) offer an alternative way to construct fragment libraries and aim to represent the associated epistemic and aleatory uncertainty. In this case, epistemic uncertainty is due to experimental errors from the determination of protein structures, while aleatory or inherent uncertainty is due to the dynamic nature, or flexibility, of proteins (Best, 2017).

Here, we present BIFROST - Bayesian Inference for FRagments Of protein STructures - a deep, generative, probabilistic model of protein backbone angles that solely uses the amino acid sequence as input. BIFROST is based on an adaptation of the Deep Markov Model (DMM) architecture (Krishnan et al., 2017) and represents the angular variables (*ϕ* and *ψ*) in a principled way using directional statistics (Mardia & Jupp, 2008). Finally, BIFROST makes it possible to evaluate the probability of a backbone conformation given an amino acid sequence, which is important for applications such as sampling the conformational space of proteins with correct statistical weights in equilibrium simulations (Boomsma et al., 2014).

## 2. Background and related work

### Probabilistic, generative models of local protein structure

Most generative, probabilistic models of local protein structure are Hidden Markov Models (HMMs) that represent structure and sequence based on the assumption of a Markovian structure (Hamelryck et al., 2012). The first such models did not include the amino acid sequence (Edgoose et al., 1998), discretised the angular variables (Bystroff et al., 2000), or used continuous, but lossy representations (Camproux et al., 1999; Hamelryck et al., 2006), making sampling of conformations with atomic detail problematic. These early models are thus probabilistic but only approximately “generative” at best. TorusDBN (Boomsma et al., 2008) was the first joint model of backbone angles and sequence that properly accounted for the continuous and angular nature of the data. Others introduced richer probabilistic models of local protein structure including Dirichlet Process mixtures of HMMs (DPM-HMMs) Lennox et al. (2010) and Conditional Random Fields (CRFs) (Zhao et al., 2010; 2008). As far as we know, BIFROST is the first deep generative model of local protein structure that aims to quantify the associated aleatory and epistemic uncertainty using an (approximate) Bayesian posterior.

### Deep Markov Models

The DMM, introduced in (Krishnan et al., 2017), is a generalisation of the variational autoencoder (VAE) (Kingma & Welling, 2014) for sequence or time series data. Related stochastic sequential neural models were reported by Fraccaro et al. (2016) and Chung et al. (2015). Published applications of DMMs include natural language processing tasks (Khurana et al., 2020), inference of time series data (Zhi-Xuan et al., 2020), and human pose forecasting (Toyer et al., 2017). Our application of the DMM and the modifications made to the standard model will be described in section 3.3.

## 3. Methods

### 3.1. Data set

BIFROST was trained on a data set of fragments derived from a set of 3733 proteins from the *cullpdb* data set (Wang & Dunbrack, 2005). Quality thresholds were (i) resolution *<* 1.6Å, (ii) R-factor *<* 0.25, and (iii) a sequence identity cutoff of 20%. For the purpose of reliable evaluation, sequences with *>* 20% identity to CASP13 targets were removed from the dataset.

Fragments containing angle-pairs in disallowed regions of the Ramachandran plot (Ramachandran et al., 1963) were removed using the Ramalyze function of the crystallography software PHENIX (Liebschner et al., 2019). The resulting data set consisted of ∼ 186000 9-mer fragments. Prior to training, the data was randomly split into train, test, and validation sets with a 60*/*20*/*20% ratio.

### 3.2. Framework

The presented model was implemented in the deep probabilistic programming language Pyro, version 1.3.0 (Bingham et al., 2019) and Pytorch version 1.4.0 (Paszke et al., 2019). Training and testing were carried out on a machine equipped with an Intel Xeon CPU E5-2630 and Tesla M10 GPU. The model trains on a single GPU and converges after 150 epochs for a total training time of approximately 34 hours.

### 3.3. Model

BIFROST consists of a DMM (Krishnan et al., 2017) with an architecture similar to an Input-Output HMM (IO-HMM) (Bengio & Frasconi, 1995). The model employs the Markovian structure of an HMM, but with continuous, as opposed to discrete, latent states (z) and with *transition and emission neural networks* instead of transition and emission matrices. Consequently, the latent states are iteratively transformed using the transition neural network, such that the value of the current latent state depends on the previous state and the (processed) amino acid information at that position (figure 2).

**Figure 2.**
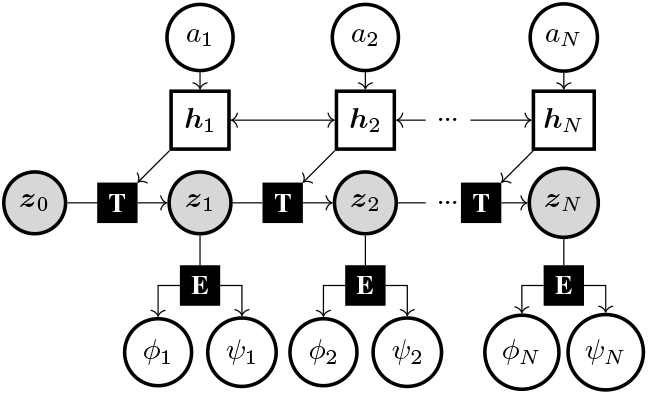
The BIFROST model. Grey nodes are latent random variables, white circular nodes are observed variables, white rectangular nodes represent hidden states from a bidirectional Recurrent Neural Network (RNN) *H*, and black squares represent neural networks. *E* and *T* denote the emitter and the transition network, respectively.

Observed angles (*ϕ* and *ψ*) are generated from the latent state sequence by applying an *emitter neural network* at each position (figure 2). Since the backbone angle *ω* is most often narrowly distributed around 180^*°*^, this degree of freedom is not included in the current version of BIFROST.

The structure of the model is shown in figure 2. For notational simplicity, the sequence of *ϕ* and *ψ* pairs will be denoted by x. The joint distribution of the latent variable z and the angles x conditioned on the amino acid sequence a with length *N* of the graphical model in figure 2 factorises as

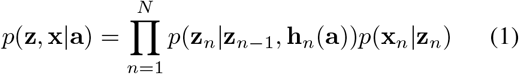

where ***h***_*n*_(a) is the deterministic hidden state generated at position *n* by a bidirectional RNN *H* with parameters ***θ***_*H*_ running across the amino acid sequence. The bidirectional RNN incorporates information from amino acids upstream and downstream of position *n*. The initial latent state z_0_ is treated as a trainable parameter and is thus shared for all sequences.

The transition densities are given by a multivariate Gaussian distribution,

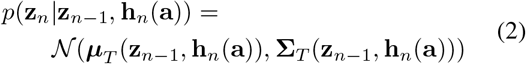

where the mean vector (***µ***_*T*_) and the (diagonal) covariance matrix (Σ _*T*_) are given by a neural network *T* parameterised by ***θ***_*T*_.

The emission densities are given by a *bivariate periodic student-T distribution* (Pewsey et al., 2007) (section 3.5) such that

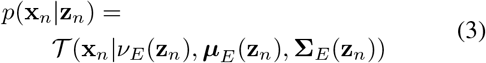

where the single, shared degree of freedom (*ν* _*E*_), the vector of two means (***µ***_*E*_), and the 2 × 2 diagonal covariance matrix (Σ _*E*_) of the distribution are given by a neural network *E* parameterised by ***θ***_*E*_.

### 3.4. Estimation

In order to perform inference of the intractable posterior, we introduce a variational distribution or *guide q* (Kingma & Welling, 2019) (figure 3), which makes use of a *combiner neural network C* parameterised by ***ζ*** _*C*_,

**Figure 3.**
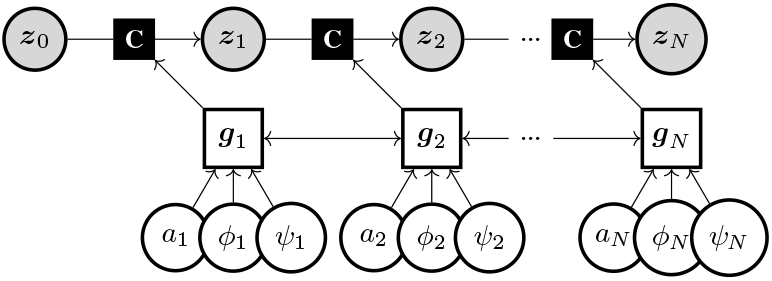
Variational distribution for approximating the posterior. Grey nodes are latent random variables, white circular nodes are observed variables, white rectangular nodes represent hidden states from a bidirectional RNN *G*, while black squares represent neural networks. *C* denotes the combiner network.

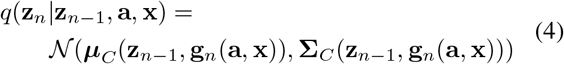

where ***g***_*n*_(***a, x***) is the deterministic hidden state generated at position *n* by a bidirectional RNN *G* with parameters ***ζ*** _*G*_ running across the amino acid sequence ***a*** and the angles ***x***.

For the parameters of the neural networks (***ζ*** _*C*_, ***ζ*** _*G*_, ***θ***_*T*_, ***θ***_*E*_, ***θ***_*H*_), point estimates are obtained using Stochastic Variational Inference (SVI), which optimises the Evidence Lower Bound (ELBO) using stochastic gradient descent (SGD) (Kingma & Welling, 2014; 2019). The ELBO variational objective is given by

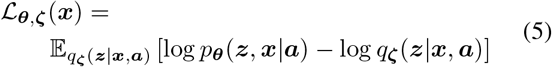

where ***ζ*** = (***ζ*** _*C*_, ***ζ*** _*G*_) and ***θ*** = (***z***_0_, ***θ***_*T*_, ***θ***_*E*_, ***θ***_*H*_) are the parameters of the guide and the model, respectively.

### 3.5. Periodic student T distribution

As angle-pairs are periodic values, i.e. distributed on a torus (Boomsma et al., 2008), they need to be modelled by an appropriate periodic distribution. Traditionally, angles are assumed distributed according to the von Mises distribution, which is defined by a mean that can be any real number and a concentration parameter, which can be any positive number. SVI showed poor performance when the von Mises distribution was used. Here, we circumvent this by representing the likelihood of the angles by a student T distribution that is wrapped around a circle (Pewsey et al., 2007). This allows for appropriate modelling of the periodicity of the angles, while being more robust with regards to outlier issues than the von Mises distribution due to the wider tails of the T distribution. It should be noted as well that Pewsey et al. (2007) showed that the wrapped student T distribution can approximate the von Mises distribution closely.

### 3.6. Neural network architecture overview

The overall architecture is based on the originally proposed DMM (Krishnan et al., 2017) with modifications. The main difference is the addition of an RNN *H* in the model that processes the amino acid sequence a, thus providing explicit conditioning on the amino acid sequence. A similar architecture was used by Fraccaro et al. (2016) for time series. In the guide, a second RNN *G* is used that processes the angles and the amino acid sequence during training. The initial values for both RNNs are treated as trainable parameters. In addition to the RNNs, the model contains an emitter network *E* and a transition network *T*, while the guide relies on a combiner network *C*.

#### Emitter architecture

The emitter network *E* parameterises the emission probabilities as stated in equation 3. *E* is a feed-forward neural network containing two initial layers that branch into three. One output branch is a single layer that outputs the degree of freedom of the Student T distribution, which is shared between the two angles. The other two branches output a mean *µ* and a standard deviation *σ* for *ϕ* and *ψ*, respectively. Each hidden layer of the neural network contained 200 neurons with rectified linear unit (ReLU) activation. Output layers for *µ* values had no activation, as the periodic distribution automatically transforms values to a range between − *π* and *π*. Output layers for *σ* and degrees of freedom *ν* used softplus activation to ensure positive, real numbered values. The architecture of *E* is depicted in figure 4.

**Figure 4.**
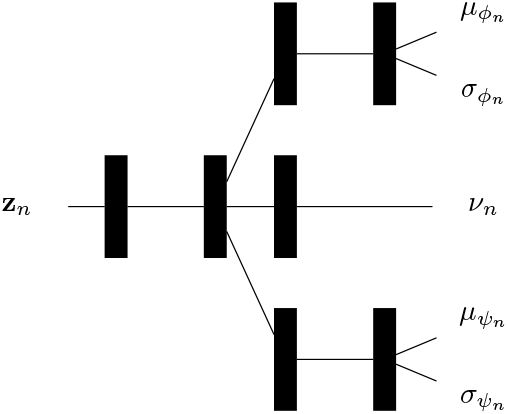
Architecture of the emitter neural network, *E*. Black rectangles represent ReLU-activated fully connected layers.

#### Transition and combiner architecture

The transition network *T* and the combiner network *C* specify the transition densities from the previous to the current latent state of the model (equation 2) and the guide (equation 4), respectively. In the original DMM (Krishnan et al., 2017), *C* was inspired by the Gated Recurrent Unit (GRU) architecture (Cho et al., 2014), while *T* was a simple feed forward network. Here, both *C* and *T* were based on GRU cells to allow for better horizontal information flow (figure 5).

**Figure 5.**
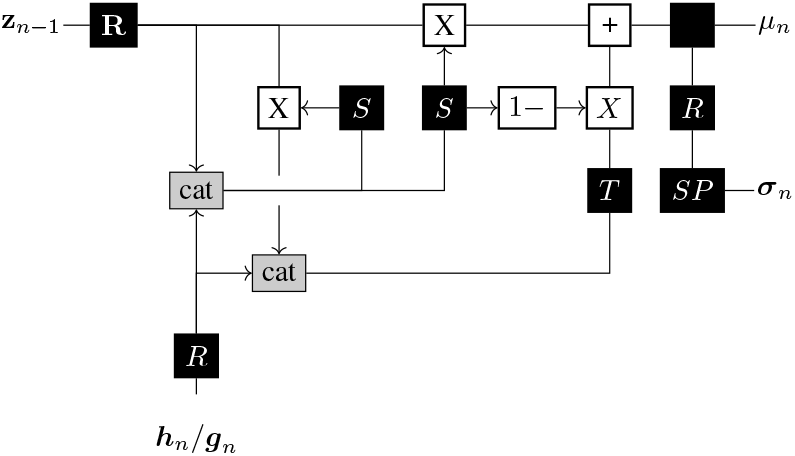
Architecture of the transition *T* and combiner *C* neural networks. Black squares represent single neural network layers activated by a ReLU (R), sigmoid (S), tanh (T), softplus (SP) or no activation. White squares represent element-wise mathematical operations. Gray squares represent tensor concatenation. Note that the network takes as input either ***h***_*n*_ or ***g***_*n*_ obtained from the RNN in the model or the guide, respectively.

The total number of parameters in BIFROST are shown in table 1.

**Table 1.** Number of parameters in BIFROST. E: Emitter, T: Transition, C: Combiner, H: model RNN, G: guide RNN, p: model, q: guide

| Neural networks | | | | | $Z_0$ | |
| --- | --- | --- | --- | --- | --- | --- |
| E | T | C | H | G | p | q |
| 24 805 | 142 280 | 142 280 | 89 200 | 90 000 | 40 | 40 |
| **Total parameters: 488 645** |  |  |  |  |  |  |

### 3.7. Hyperparameter optimization

A simple hyperparameter search was performed with the test ELBO as the selection criterion (data not shown). The final model was trained with a learning rate of 0.0003 with a scheduler reducing the learning rate by 90% when no improvement was seen for 10 epochs. Minibatch size was 200. The Adam optimiser was used with a *β*_1_ and *β*_2_ of 0.96 and 0.999 respectively. The latent space dimensionality was 40. All hidden activations (if not specified above) were ReLU activations. We employed norm scaling of the gradient to a norm of 10.0. Finally, early stopping was employed with a patience of 50 epochs.

### 3.8. Sampling from the model

The BIFROST model (figure 2) is designed with explicit conditioning on amino acid sequences allowing a simple and efficient ancestral sampling approach that eliminates the need for using the guide for predictions. Thus, the guide is used solely for the purpose of model estimation and is discarded upon sampling.

### 3.9. Fragment library generation and benchmarking

Fragment libraries are a collection of fragments, consisting of typically 3 or 9 amino acids with known backbone angles. Here, we focus on fragments of nine amino acids. For each fragment in a protein, 200 possible backbone conformations are sampled from BIFROST resulting in a set of (*L −* 8) × 200 fragment candidates, where *L* is the number of amino acids in the protein. These candidates are compared to the observed fragment by calculating the angular root mean square deviation (RMSD) between the corresponding angles as proposed in Boomsma et al. (2008). The choice of 9-mer fragments and the 200 samples per fragment were made to emulate the default behavior of the Rosetta fragment picker (see below), for fair comparison.

The aggregated quality of fragment libraries are generally represented by two metrics; *precision* and *coverage*. Precision is defined as the fraction of candidates with an RMSD to the observed below a certain threshold, whereas coverage is the fraction of positions covered by at least one candidate with an RMSD below a certain threshold. Evaluating the precision and coverage at increasing thresholds yields two curves, and the quality of the fragment library is quantified by the area under these two curves.

BIFROST was benchmarked against Rosetta’s fragment picker (Gront et al., 2011) using the precision and coverage metrics. The fragment picker was run using default parameters, picking 200 fragments per position. Secondary structure predictions were performed using SAM-T08 (Karplus, 2009), PSIPRED (Jones, 1999) and Jufo (Leman et al., 2013). Sequences that were homologous to the targets were excluded (*–nohoms* flag).

Fragment libraries were generated for all available regular (denoted “T”) targets from the latest installment of the bi-annual protein structure prediction competition Critical Assessment of Techniques for Protein Structure Prediction (CASP13).

### 3.10. Runtime comparison

In order to compare the runtime of BIFROST to that of the fragment picker, nine proteins of varying lengths were selected. Both tools generated 200 samples per fragment. The experiment was run on the same 32-core machine for both the fragment picker and BIFROST.

## 4. Results

To show that the model is able to capture general protein backbone behavior, angles were generated conditioned on the sequences of 5000 previously unseen fragments and compared to the observed angles. The model was able to recreate the observed Ramachandran plots with minimal added noise (figure 6).

**Figure 6.**
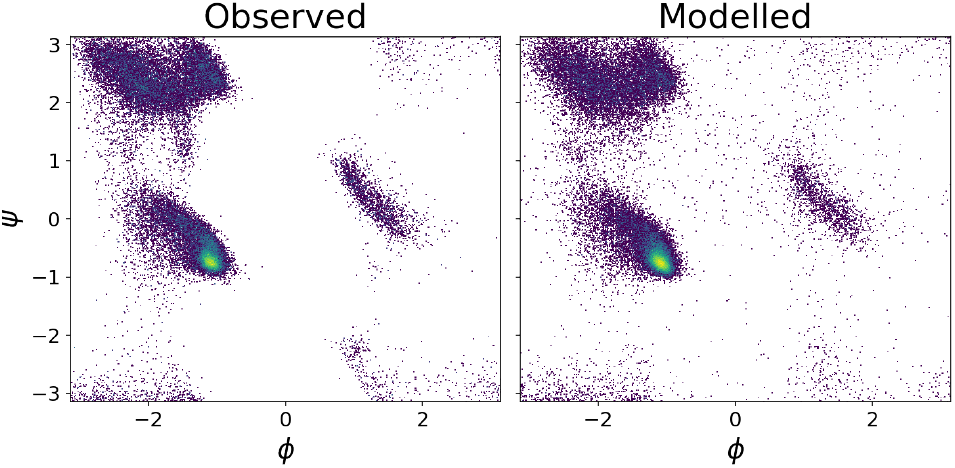
Observed and modelled aggregated Ramachandran plots

While most amino acids show angle distributions similar to the background in figure 6, glycine and proline are exceptions due to the nature of their side chains. The side chain of glycine is a single hydrogen atom, allowing the backbone to be exceptionally flexible, while the side chain of proline is covalently linked to the backbone restraining the conformational space. The modelled distribution of angles for these two unique cases, along with leucine to represent the general case, show that the model is able to capture specific amino acid properties (figure 7).

**Figure 7.**
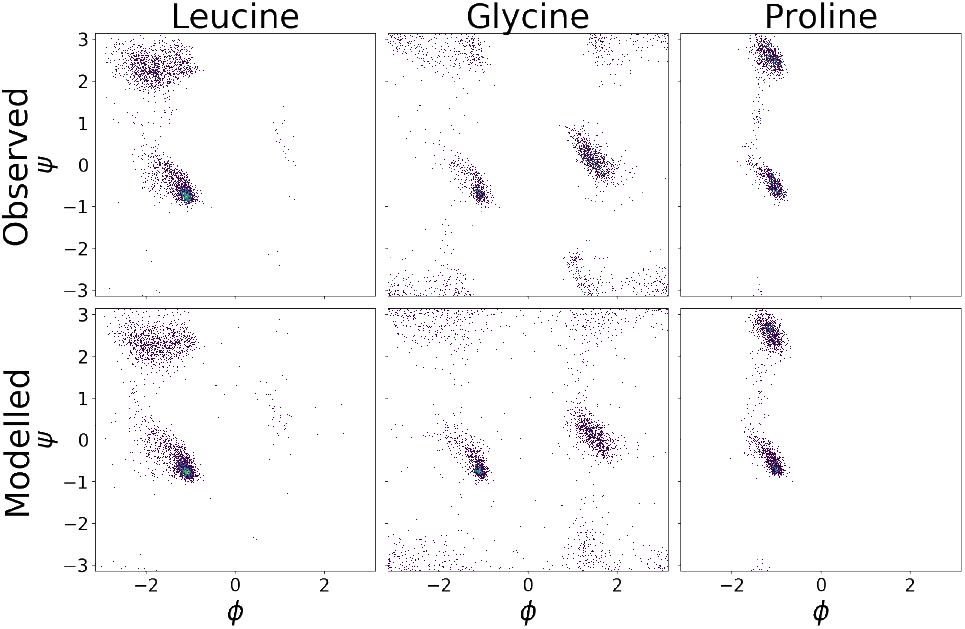
Amino acid specific Ramachandran plots

The left side of figure 8 shows a thin, smoothed coil representation of 100 samples from BIFROST conditioned on example 9-mer fragments that were observed to be either *α*-helix, *β*-strand, or coiled. The right side shows distributions of backbone RMSDs of 5000 sampled fragments to the observed structure from BIFROST and as picked by Rosetta’s fragment picker.

**Figure 8.**
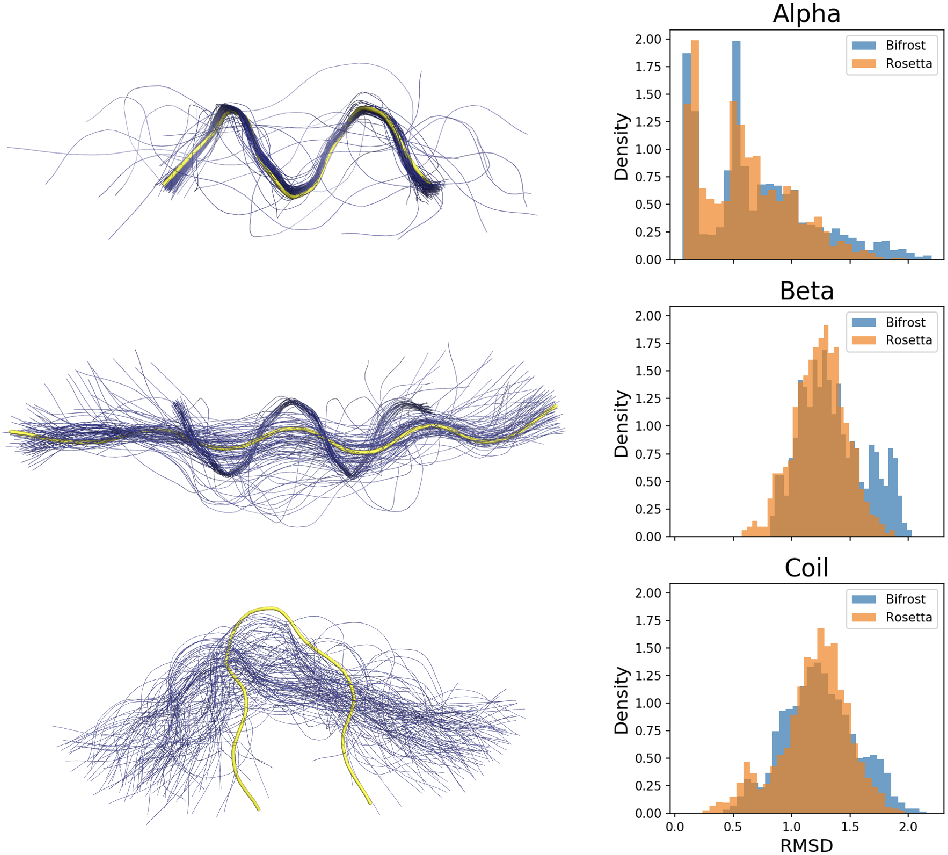
Left: 100 samples of backbone dihedral angles (blue) superimposed on the observed structures (yellow). For clarity, the backbones are represented as thin, smoothed coils instead of traditional cartoon representations. Right: Aggregated RMSDs of BIFROST-sampled conformations and conformations picked by Rosetta’s fragment picker for sequences observed as *α*-helix, *β*-strand, and coil respectively.

The RMSDs were generally distributed towards 0Å for the *α*-helix case, showcasing BIFROSTs ability to predict this well defined secondary structure element. The model has more difficulty modelling *β*-strands and coils. However, the distributions of the RMSDs are nearly identical to those produced by the fragment picker. For coil fragments, the RMSDs were distributed around 3Å reflecting the inherent variability of those fragments.

BIFROST was benchmarked against Rosetta’s fragment picker (Gront et al., 2011) on all publicly available CASP13 regular targets. BIFROST generated fragment libraries with comparable precision and coverage to the fragment picker (figure 9).

**Figure 9.**
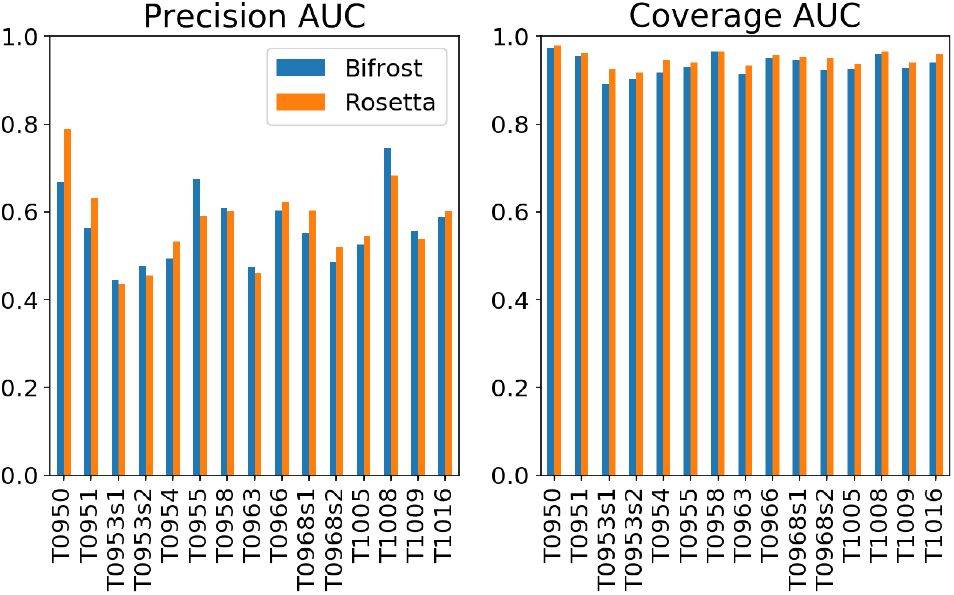
Comparison of fragment libraries generated by BIFROST, relying on just the amino acid sequences, against Rosetta’s fragment picker, which uses external information and relies on ensemble predictions of secondary structure.

Finally, BIFROST enables efficient sampling of fragment libraries. The runtime of BIFROST and the fragment picker are compared in figure 10 on a set of nine proteins of varying lengths. Both runtimes roughly scale linearly with protein length, but BIFROST has a smaller constant term than the fragment picker.

**Figure 10.**
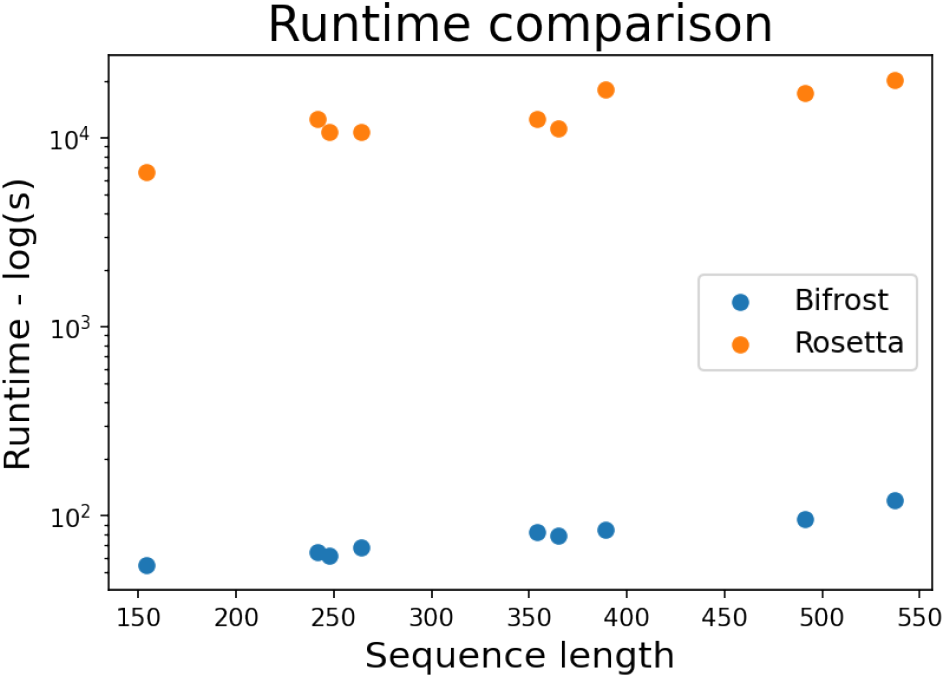
Runtime comparison between Rosetta’s fragment picker and BIFROST on a set of nine proteins of varying lengths.

## 5. Discussion

BIFROST is a deep, generative model of local protein structure conditioned on sequence that provides a probabilistic approach to generating fragment libraries.

The quality of the generated fragment libraries is on par with Rosetta’s fragment picker, despite using much less information, such as an ensemble of secondary structure predictors. Due to the probabilistic nature of BIFROST, distributions tend to be slightly wider than those resulting from picking structural fragments from the PDB based on sequence similarity. This wider distribution plausibly reflects the dynamic nature of protein structure, which is not captured in the experimental data provided by static X-ray structures.

The model was estimated using SVI, relying on the ELBO variational objective. As the ELBO provides a lower bound on the log evidence (Kingma & Welling, 2014), we can evaluate the probability of a specific local structure given the sequence, simply by evaluating the ELBO. Evaluating the probability of fragments is crucial for correct sampling of the conformational space, for example in the case of equilibrium simulations of protein dynamics (Boomsma et al., 2014). The probabilities assigned by BIFROST can be used to decide how often a fragment should be sampled in the folding process. In contrast, existing methods do not provide an explicit measure of fragment confidence.

In this paper the focus was kept on fragments of nine residues for ease of comparison to the fragment picker. However, the DMM architecture of BIFROST allows generation of fragments of arbitrary length but with an observed drop-off in performance as the length of fragments are increased (data not shown).

Existing methods rely heavily on the availability of multiple sequence alignments (MSA) and other information, such as secondary structure predictors. As MSAs are not available for orphan proteins or synthetic proteins, the need for pure sequence based models is evident.

## 6. Acknowledgements

We acknowledge funding from the Innovation Fund Denmark under the grant “Accelerating vaccine development through a deep learning and probabilistic programming approach to protein structure prediction”. We thank Wouter Boomsma for help with the wrapped student T distribution

